# Deciphering human heart failure with preserved ejection fraction (HFpEF) at single cell resolution

**DOI:** 10.1101/2025.04.02.646923

**Authors:** Lukas Zanders, Simone F Glaser, Mariano Ruz Jurado, Moritz Brandt, David John, Luka Nicin, David Rodriguez Morales, Wesley T Abplanalp, Tara Procida-Kowalski, Franziska Ganß, Evelyn Ullrich, Alisa Debes, Penelope Pennoyer, Marek Bartkuhn, Reinhard B Dettmeyer, Andreas M Zeiher, Eike Nagel, Philip Wenzel, Stefanie Dimmeler

## Abstract

**BACKGROUND:** Heart failure with preserved ejection fraction (HFpEF) is a complex and growing condition, representing over half of all heart failure cases. Despite its high morbidity and mortality, its heterogeneity and limited therapeutic options pose significant challenges. Understanding the molecular mechanisms driving HFpEF is essential for the development of new therapies to improve patient outcomes.

**METHODS:** We performed single-nucleus RNA sequencing of nuclei obtained from endomyocardial biopsies of six patients with HFpEF. The obtained dataset was integrated with a dataset of 12 healthy human hearts and their transcriptomic differences were analyzed.

**RESULTS:** After quality control and integration of the datasets, nine major cardiac cell types were annotated. HFpEF cardiomyocytes were characterized by a reduction in genes associated with aerobic respiration and fatty acid metabolism and showed an upregulation of RHOA/ROCK1 signaling, which was validated using immunofluorescence staining in human HFpEF myocardial sections. Endothelial cells exhibited signs of increased apoptosis, SEMA3 signaling and signs of reduced VEGFA signaling as well as a reactivation of a fetal gene signature. In line with a prominent role of cardiac fibrosis in HFpEF, we observed increased signs of fibroblast activation and proliferation, and reduced signs of IFNɣ signaling in HFpEF which was most pronounced in activated fibroblasts. Treatment of human cardiac fibroblast with rhIFNɣ resulted in decreased collagen contents. Macrophages from HFpEF myocardium showed a pro-inflammatory transcriptomic signature and showed increased expression of MHC-II molecules. This was associated with signs of an increased IFNɣ response.

**CONCLUSION:** Our results provide insights into the transcriptional diversity of HFpEF recapitulating structural, functional, and molecular hallmarks of the disease and provide mechanistic insights which might represent therapeutic targets and biomarkers to improve outcome of patients with HFpEF.

**CLINICAL PERSPECTIVE:** *What is new?:*

- We provide a single-nucleus RNA sequencing (snRNA-Seq) dataset from human HFpEF myocardium and demonstrate feasibility of snRNA-Seq from endomyocardial biopsies
- The snRNA-Seq data confirms signs of known molecular hallmarks of HFpEF, such as metabolic changes, inflammation and fibrosis
- We identify signs of regulating cellular mechanisms underlying these hallmarks, such as cytoskeleton remodeling via RhoA/ROCK1 in cardiomyocytes, and differential interferon gamma signaling in stromal and immune cells

*What are the clinical implications?:*

- We provide several cell type-specific cellular mechanisms which might serve as biomarkers or therapeutic targets in the treatment of HFpEF

## INTRODUCTION

Heart failure with preserved ejection fraction (HFpEF) is a complex and increasingly prevalent condition, accounting for over half of all heart failure cases. Despite its high morbidity and mortality, HFpEF remains a major challenge in clinical practice, partially due to its heterogeneity in underlying myocardial disease, comorbidities, and disease mechanisms [1–3]. The complexity results in the lack of effective targeted therapies for most etiologies. While new therapies, especially SGLT2 inhibitors, can reduce heart failure hospitalizations, there are no therapies available which reduce mortality rates in HFpEF patients [4]. In some instances, such as in cardiac transthyretin amyloidosis, investigation of molecular mechanisms driving disease progression have led to the development of effective therapies and in combination with the development of readily available diagnostic tests, reduced morbidity and mortality, demonstrating that a thorough understanding of disease mechanisms can improve patients’ outcomes [5, 6]. However, for most etiologies of HFpEF, the underlying molecular disease-associated mechanisms remain largely elusive. Importantly, while previously tested medical therapies primarily target hormonal and hemodynamic consequences of heart failure, no therapies for cellular and molecular mechanisms driving disease progression are currently available.

Several clinical risk factors, such as obesity, increased age, diabetes mellitus, and hypertension have been associated with the development of HFpEF and pose the main target of therapeutic options. Studying the molecular mechanisms driving HFpEF is critical to better understand its pathophysiology in order to deliver novel therapeutic approaches and improve patient outcomes [1]. Unlike heart failure with reduced ejection fraction, HFpEF lacks a single dominant pathophysiologic mechanism, reflecting its heterogeneity in humans [3]. Existing animal models attempt to replicate key features of HFpEF, such as diastolic dysfunction, left ventricular hypertrophy, and systemic inflammation. However, they do not fully recapitulate the complex interactions often induced by combinations of predisposing factors and comorbidities [3, 7]. Moreover, human myocardial tissue samples are sparse as HFpEF patients rarely undergo procedures during which large amounts of tissue are easily obtainable, such as heart transplant. The combination of a lack of animal models fully recapitulating human disease and the sparsity of human tissue samples aggravate the investigation of pathophysiologic mechanisms.

Few cellular and molecular disease mechanisms have previously been shown to play major roles in the development and progression of HFpEF. For example, metabolic changes with a decrease in fatty acid oxidation and aerobic respiration, as well as signs of reduced glycolysis were shown as hallmarks of HFpEF [2, 8, 9]. Microvascular dysfunction with capillary rarefaction and endothelial dysfunction results in relative myocardial hypoxia and dysfunction [10–12]. As diastolic dysfunction is a key feature of HFpEF, the underlying mechanisms have been extensively studied and point towards a role of activated fibroblasts and extracellular matrix (ECM) deposition [13–15]. Interestingly, not only interstitial fibrosis, but also increased cardiomyocyte stiffness and hence, dysfunctional relaxation may also contribute to diastolic dysfunction in HFpEF [16]. Importantly, low-grade inflammation with contributions from the humoral and adaptive immune system are key features of HFpEF [17]. For example, the abundance of macrophages is increased in the myocardium from HFpEF patients [15]. This is accompanied by increased circulating levels of pro-inflammatory cytokines such as IL-6, which predisposes for the development of new-onset HFpEF [18].

However, the regulating factors of cellular disease mechanisms, which might represent therapeutic targets, as well as many molecular pathophysiologic contributors to HFpEF remain poorly understood. For the development of targeted and effective treatments, it is therefore essential to better understand these mechanisms and upstream regulators with high cellular resolution.

Single-cell technologies have been the basis for the development of in-depth insights into human pathologies by generating human cell atlases, including – for example - single-nuclei profiling of the healthy human heart [19], hereditary cardiomyopathies [20] human dilated and hypertrophic cardiomyopathy [21, 22]. However, information on the transcriptional patterns in HFpEF are sparse particularly given the limitation for tissue samples, which relies on endomyocardial biopsies [23]. Here we used tissue biopsies of patients with HFpEF to gain insights into the transcriptional changes at single-cell resolution.

## METHODS

### Patients

Endomyocardial specimens were acquired from five patients undergoing transcatheter endomyocardial biopsy (EMB) for diagnostic workup for HFpEF at the University Clinic Mainz, Germany. The usage of tissue samples for research purposes was approved by the institutional ethics review board of the University Mainz (application number 837.337.15 (10105)). One sample was obtained from the DECIPHER HFpEF clinical trial which was approved by the ethics review board of the medical faculty of the Goethe-University Frankfurt (application number 273/17). All patients gave informed written consent and the trial was performed in compliance with institutional guidelines and the Declaration of Helsinki.

Samples were obtained from the left ventricle in five patients and from the interventricular septum from right ventricle in one patient. HFpEF was defined as the combination of symptoms of heart failure (dyspnea, fatigue, peripheral edema), a left ventricular ejection fraction of ≥50% and echocardiographical or histological signs of structural heart disease, according to the current guidelines by the American Heart Association, American College of Cardiology, Heart Failure Society of America and the European Society for Cardiology [24, 25].

Patient’s baseline characteristics were compared using the Wilcoxon rank sum test for continuous variables and Fisher’s exact test for categorial variables.

### Nuclei isolation, snRNA-seq library preparation and sequencing

EMBs were snap-frozen in liquid nitrogen immediately after acquiring the samples and stored at −80°C until further processing. Nuclei were isolated as previously described [22]. Briefly, two to three EMB, each weighing about 3 mg, per patient were manually minced on ice and transferred to a hypoosmolar lysis buffer containing Ambion RNase inhibitor (final concentration 0.6 U/µL; ThermoFisher AM2684). EMBs were homogenized using a Dounce homogenizer. After homogenization, samples were filtered using 40 µm (Corning CLS431750) and 20 µm (PluriSelect 43-50020) cell strainers. Nuclei were stained with 7-Aminoactinomycin D (Merck SML1633) for 20 minutes on ice and sorted using a FACSAria Fusion 6B-3R-2UV sorter (BD Bioscience) to separate nuclei from debris. Nuclei were subsequently washed and visually checked for integrity. If nuclei were intact, we proceeded to library construction.

Libraries for snRNA-Seq were constructed with the 10X Genomics Chromium 3’ v3.1 platform, using a Chromium controller and Chromium Single Cell 3’ v3 reagents according to the manufacturer’s instructions. 800 – 3000 nuclei per sample were loaded per Chip. Quality control of the libraries was performed using an Agilent Tapestation 4200 before sequencing on an Illumina NovaSeq 6000 with a targeted sequencing depth of 50,000 reads per nucleus.

### Sequencing data processing

The sequencing reads were processed and mapped using the CellRanger single cell software suite (v7.0.0, 10X Genomics) utilizing the human reference genome GRCh38-2020, including intronic reads. Count matrices were processed using CellBender (v0.3.2) [26] to remove ambient RNAs. All subsequent analyses were performed in R (v4.4.0) and Seurat (v5.0.2) [27]. For quality control, nuclei with <300 or >6000 features or mitochondrial counts > 5% were removed from the dataset. Count normalization was performed using the default settings from Seurat.

We integrated the snRNA-Seq data with data from septal myocardium from twelve healthy donors [19]. Dimensionality reduction was performed over highly variable genes using principal component analysis (PCA). Canonical Correlation Analysis (CCA) was performed on the PCA embedding to correct for batch effects. The CCA dimension reduction was used for subsequent Louvain clustering (resolution 0.3). Cell type annotation was performed with the CellTypist (v1.6.2) package facilitating the reference Healthy_Adult_Heart model v1 [28]. Validation of annotation was performed using canonical cell type markers. Visualization of the clustering was done using Uniform Manifold Approximation and Projection (UMAP). For comparison of overall transcriptional differences between samples, the average normalized unique molecular identifier (UMI) counts for each gene were calculated per sample. The pseudo-bulk expression matrix was used for Pearson correlation analysis and the correlation coefficients were plotted using corrplot v0.92.

### Cluster distribution and differential gene expression analyses

Distributions of annotated cell types were calculated after down-sampling of the whole dataset to the smallest number of nuclei per condition and cell type to achieve similar numbers of overall nuclei. Significance testing of differential distribution was tested using a Chi-square test and if the p value was < 0.05, a Wilcoxon Rank-sum test with Benjamini-Hochberg p value adjustment was performed on the distribution data per sample. Cell type distributions were visualized with stacked bar graphs using ggplot2.

Differentially Expressed Genes (DEG) were calculated for each cell type using the Model-based Analysis of Single-cell Transcriptomics (MAST) with Bonferroni correction [29]. As MAST is limited with regards to the analysis of very low read counts, some analyses were performed using the Wilcoxon Rank-sum test with Bonferroni correction. The analysis in which the latter test was used are indicated within the figure legends. Genes were considered differentially expressed at an adjusted p value < 0.05. For the comparative analysis of the number of DEGs per cell type, we down-sampled the highly abundant cell types to the smallest number of nuclei per condition and cell type (150 nuclei) and performed differential gene expression (DGE) analyses using MAST. All DEGs, irrespective of log2 fold change were plotted in the respective bar graph. For all other DGE analyses, the whole dataset was analyzed. Venn Diagrams were created using the Venn webtool of the Bioinformatics & Evolutionary Genomics Department of the University Gent (https://bioinformatics.psb.ugent.be/webtools/Venn/).

### Functional enrichment, gene-set annotation and visualization of gene expression

For downstream analyses, all ribosomal RNAs (rRNA) and mitochondrial genes were removed from the DGE analysis outputs, as their abundance is strongly influenced by tissue integrity and the efficacy of rRNA depletion during snRNA-Seq library construction. All DEGs with an adjusted p value of < 0.05 and an average log2 fold change of < −1 for down-regulated and > +1 for up-regulated genes were subjected to functional enrichment analyses using the metascape platform with the ‘Express Analysis’ [30]. The output files were used for the visualization using ggplot2 (v3.5.1). The summary terms were plotted sorted by the ascending p value. For visualization of gene expression via bubble, bar and boxplots, the DOtools package (v0.0.1) was used.

For visualization of gene expression in heatmaps, the average normalized counts of the respective genes were calculated, and Z-scores were calculated for rows and columns. The heatmaps were created using ComplexHeatmap (v2.22.0). For signaling pathway visualization, all DEGs of the respective cell type with an adjusted p-value of < 0.05 as determined by MAST were used as input for pathway illustration using pathview (v1.46.0).

### Ligand receptor interaction analysis

Ligand receptor analysis were performed through CellChat (v2.1.2) according to the standard workflow as recommended by the authors [31]. For control and HFpEF datasets, independent CellChat objects were created and subsequently merged to facilitate comparative analyses. As the cell type populations differed in size, we used down-sampled datasets to avoid the overrepresentation of abundant cell types. As the majority of predicted ligand-receptor interactions for endothelial cells were Collagen, Laminin, Ptprm and Pecam1 and blunted other predictions, they were removed from visualization using netAnalysis_signalingChanges_scatter().

### Subclustering and re-annotation

We identified sub-populations of fibroblasts and macrophages by re-clustering using the Louvain algorithm and annotating with CellTypist. Two sub-populations of fibroblasts were identified: activated and non-activated fibroblasts, which was confirmed using canonical activation markers (COL1A1, COL1A2, COL3A1, POSTN and ACTA2). Two sub-populations of macrophages were identified: tissue resident and bone-marrow-derived macrophages, which was confirmed using markers (LYVE1, FOLR2, CD163L1, CCR2, S100A4, ADGRE5). Quantification and study of the distribution was performed on a down-sampled dataset as described above.

### Gene-set based scoring

Cell cycle prediction in fibroblasts was performed as previously described [32]. This approach facilitates marker gene sets for cell cycle stages and uses their aggregate average expression after subtraction of the aggregate average expression of a random control gene set. Quantification of predicted cell cycle stages was performed on a downsampled dataset as described above, which was similarly performed for all distribution analyses described below.

To calculate interferon gamma response and production scores, we used published marker gene sets [33] in the AddModuleScore() function, facilitating a similar approach as described above.

### Bulk RNA-sequencing analysis

Publicly available datasets from human HFpEF and control myocardial samples (10.5281/zenodo.4114617) [23] were downloaded from the zenodo platform, and human healthy fetal and adult myocardium (GSE126569) [34] were downloaded from the GEO database. We used DESeq2 for calculation of DEGs (v1.44.0). We used the Wald test for DEG calculation with Benjamini-Hochberg post-hoc adjustment and used the default analysis pipeline of DESeq2.

### Immunofluorescence stainings, imaging and quantification

Endomyocardial biopsies from the DECIPHER HFpEF trial and myocardial samples obtained from controls without signs of heart disease post-mortem (Forensic Medicine, University Giessen) were fixed in 4% paraformaldehyde over night at 4°C, embedded in paraffin and stored at room temperature until further processing. 4 µm sections were cut and processed for immunofluorescence staining. Deparaffinization was performed by heating the slides to 60°C for 30 minutes followed by 2× 10 minutes in RotiHistol (Carl Roth 6640.1) and dehydration. Antigen retrieval was performed in 0.01M citrate buffer at pH 6.0 in a pressure cooker for 90 seconds. After permeabilization with 0.2% Triton X-100 (ThermoFisher T8787), sections were incubated in a blocking solution for one hour at room temperature (2% Donkey Serum (abcam ab7475), 3% BSA (Sigma-Aldrich A1595), 0.2% Triton X-100 in DPBS (130225 PAA Laboratories)) and incubated with a ROCK1 antibody (abcam Alexa Fluor® 488 Anti-ROCK1 [EPR638Y], 1:50) and Ulex Europaeus Agglutinin I (Vector Laboratories B-1065-2, 1:100) in blocking solution at 4°C, overnight. After washing, streptavidin-conjugated Alexa Fluor® 555 (ThermoFisher S21381, 1:200) was added, slides were washed and embedded in Fluoromount-G (Invitrogen 00-4958-0) with Hoechst 33342 (Life Technologies H3570, 1300). Imaging was performed on a Leica STELLARIS confocal microscope and signal quantification was done using Volocity v7 (Quorum Technologies).

### Treatment of human cardiac fibroblasts with IFNɣ, immunofluorescence staining, imaging and quantification

Primary human cardiac fibroblasts (Promocell, C-12375, LOT 495037.1) were cultured in Fibroblast Growth Medium 3 (Promocell C-23025) and seeded into 18-well chambered cell culture coverslips (ibidi 81816). After 24 hours, cells were treated with the indicated concentrations of recombinant human interferon gamma (R&D Systems, 285-IF) for 72 hours. Cells were stained as previously described [35]. Briefly, cells were fixed in 4% paraformaldehyde for 10 minutes at room temperature, permeabilized in 0.1% Triton X-100 for 15 minutes and blocked in 5% donkey serum. Staining with rabbit anti-COL1A1 (Cell Signaling 72026, 1:100) and phalloidin Oregon Green 488 (ThermoFisher O7466, 1:100) was performed at 4°C overnight. After washing, secondary antibody (donkey anti-rabbit-647, ThermoFisher A-31573, 1:200) was added for one hour at room temperature. Cells were washed and mounted with Fluoromount-G (Invitrogen 00-4958-0).

Imaging was performed on a Leica STELLARIS confocal microscope and signal quantification was done using Volocity v7 (Quorum Technologies).

### Statistical testing of imaging quantification data

Normalized Volocity quantification results were analyzed using GraphPad Prism v10.1.1. For in vitro data, a Kruskal-Wallis-Test with Dunn’s post-hoc testing for pairwise comparison was performed. For immunofluorescence quantification data from human sections, a Mann-Whitney U test was used. Data are presented as mean + SEM and the p values are indicated.

### Data Availability

Single nuclei RNA sequencing datasets will be made publicly available from ArrayExpress upon publication of the manuscript.

### Code Availability

Before publication, the code used for the analysis can be accessed through https://figshare.com/s/1d4a75fc3e69112fdc08 and will be made publicly available on GitHub (https://github.com/lukas-zanders/2025_Zanders_HFpEF) upon publication.

## RESULTS

### Cellular landscape in HFpEF

We processed and generated single-nucleus RNA sequencing data of endomyocardial biopsies deriving from six patients with HFpEF (**Fig. 1a**) as defined by symptoms of heart failure, a left ventricular ejection fraction of ≥ 50% and echocardiographical or histological signs of structural heart disease, according to current guidelines by the European Society for Cardiology and the American Heart Association [24, 25]. HFpEF patients exhibited more cardiovascular risk factors, such as hypertension and diabetes. The etiology of HFpEF remained unclear in four cases, while one patient had transthyretin amyloidosis and one patient had hypertrophic cardiomyopathy, reflecting the heterogeneous background of HFpEF. (**Fig. 1b**; for detailed patient characteristics see **Table 1**).

**Figure 1.**
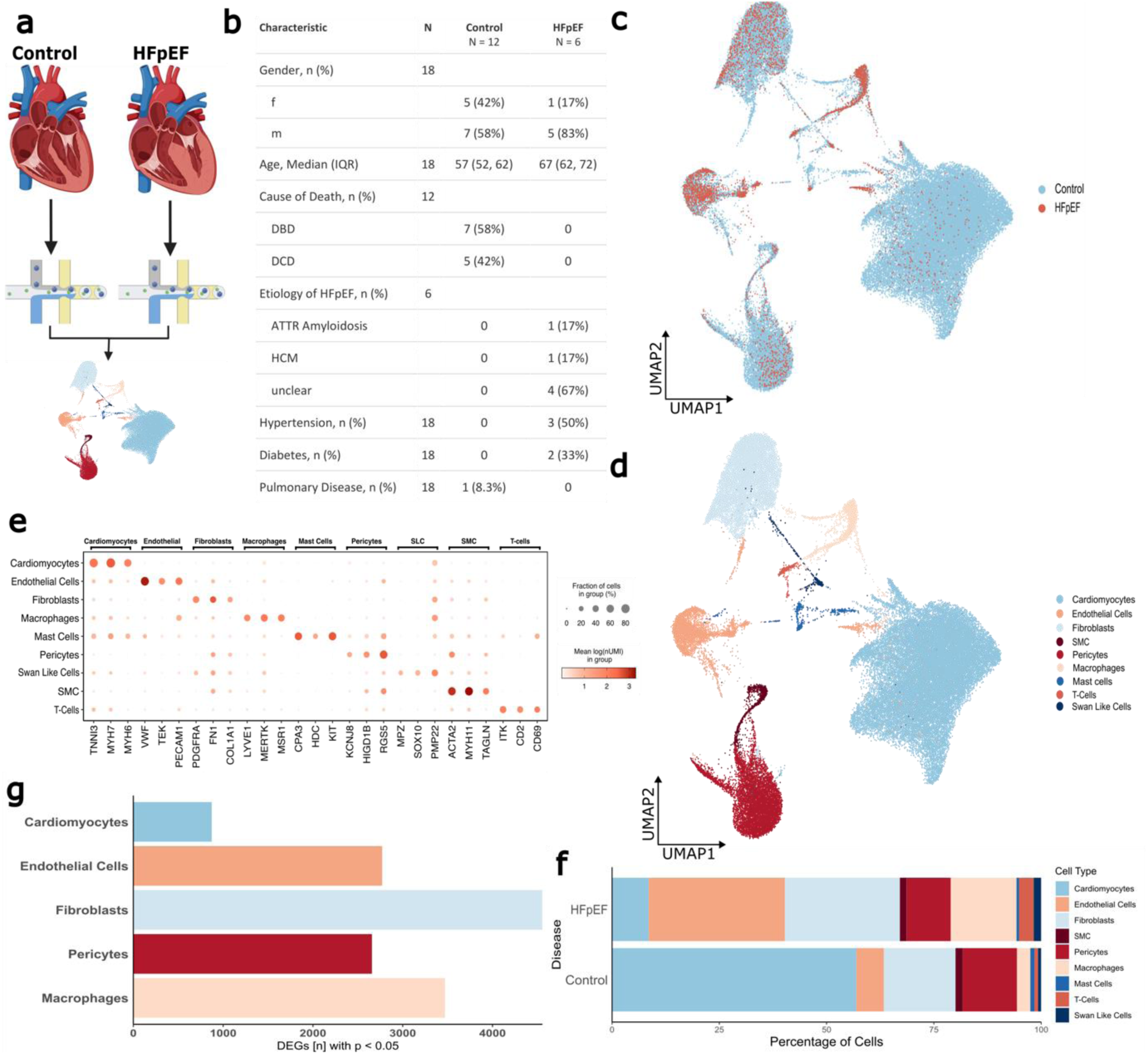
An atlas of the human HFpEF heart. **a**, Schematic illustration of the workflow. Nuclei from HFpEF myocardium were isolated, subjected to snRNA-Seq and combined with a publicly available control dataset for downstream analyses. Illustration created with biorender.com **b**, Summary of the baseline characteristics of the Control and HFpEF patients included in the trial. **c**, UMAP over all samples showing the distribution of Control and HFpEF nuclei. **d**, UMAP over all samples (60,199 nuclei) passing QC showing the cell type annotation. **e**, Average expression of canonical cell type marker genes across the cell types. **f**, Number of DEGs using MAST in a subsampled dataset of the most abundant cell types. **g**, Cell type distribution in HFpEF and Control groups as percentage of the respective group. HFpEF, Heart Failure with preserved Ejection Fraction; snRNA-Seq, single nucleus RNA sequencing; UMAP, Uniform Manifold Approximation and Projection, QC, Quality Control; DEGs, Differentially Expressed Genes; MAST, Model-based Analysis of Single-cell Transcriptomics.

**Table 1.**
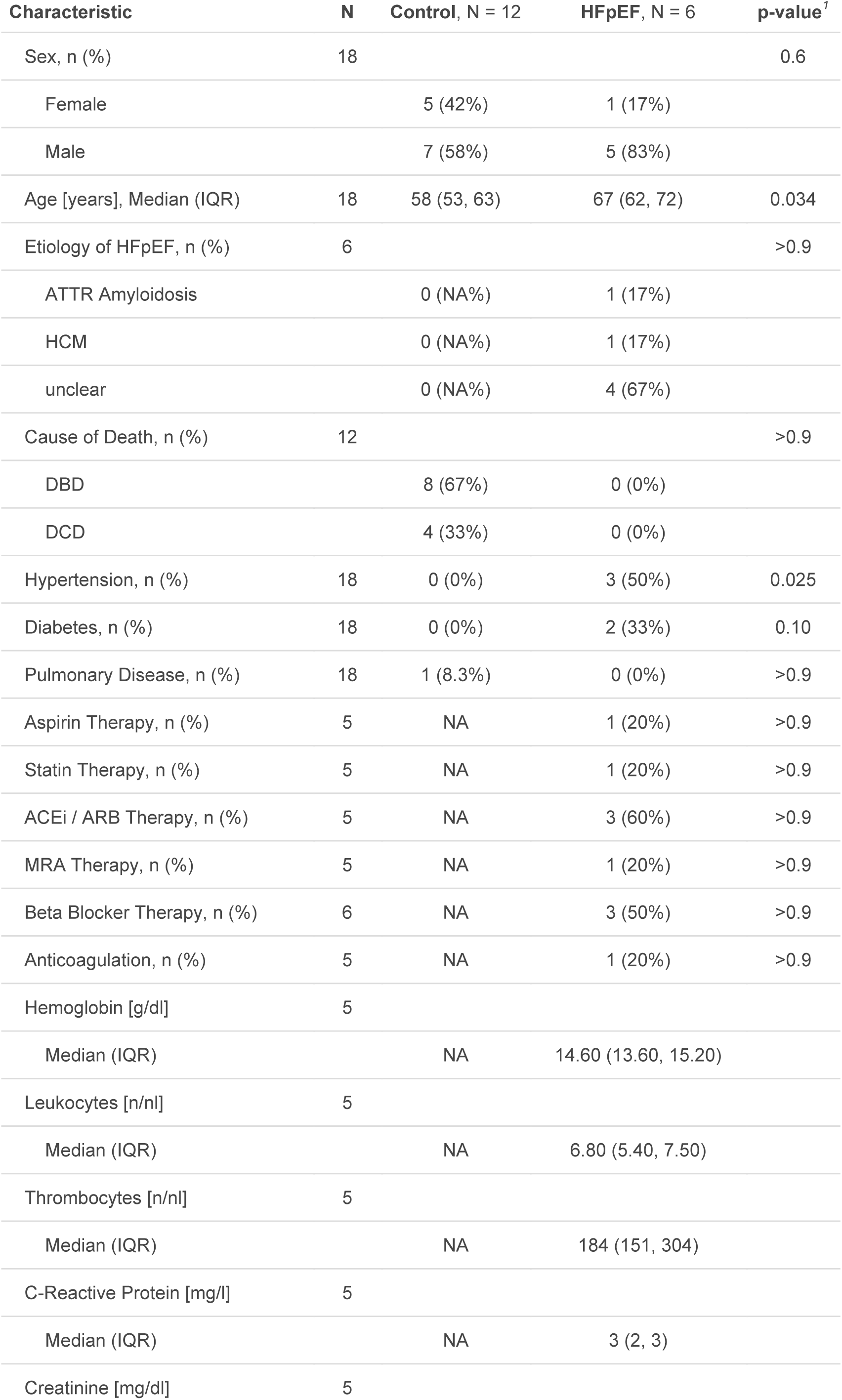
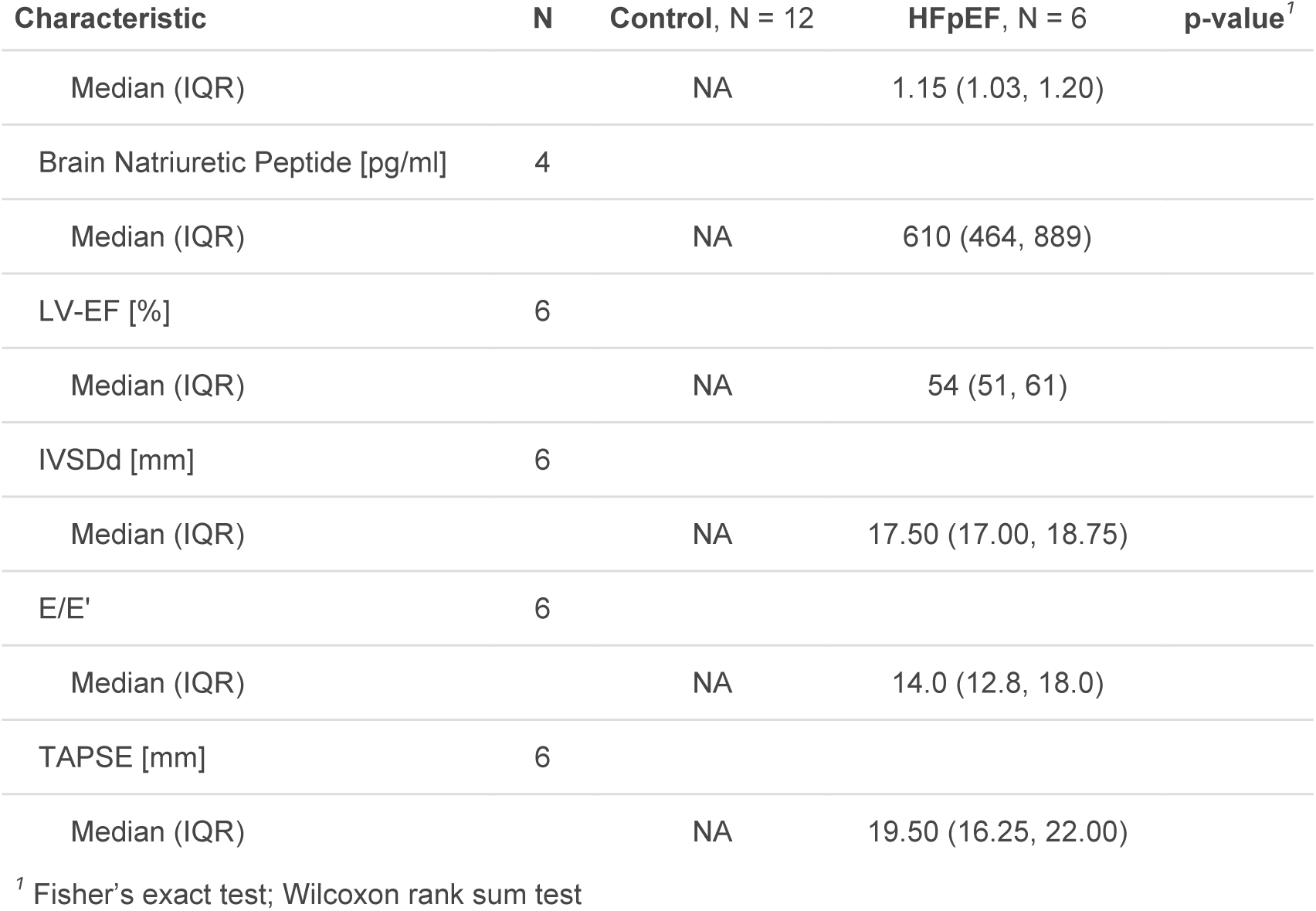
Patient’s and donor’s baseline characteristics.

| Characteristic | N | Control, N = 12 | HFpEF, N = 6 | p-value<sup>†</sup> |
| --- | --- | --- | --- | --- |
| Sex, n (%) | 18 |  |  | 0.6 |
| Female |  | 5 (42%) | 1 (17%) |  |
| Male |  | 7 (58%) | 5 (83%) |  |
| Age [years], Median (IQR) | 18 | 58 (53, 63) | 67 (62, 72) | 0.034 |
| Etiology of HFpEF, n (%) | 6 |  |  | >0.9 |
| ATTR Amyloidosis |  | 0 (NA%) | 1 (17%) |  |
| HCM |  | 0 (NA%) | 1 (17%) |  |
| unclear |  | 0 (NA%) | 4 (67%) |  |
| Cause of Death, n (%) | 12 |  |  | >0.9 |
| DBD |  | 8 (67%) | 0 (0%) |  |
| DCD |  | 4 (33%) | 0 (0%) |  |
| Hypertension, n (%) | 18 | 0 (0%) | 3 (50%) | 0.025 |
| Diabetes, n (%) | 18 | 0 (0%) | 2 (33%) | 0.10 |
| Pulmonary Disease, n (%) | 18 | 1 (8.3%) | 0 (0%) | >0.9 |
| Aspirin Therapy, n (%) | 5 | NA | 1 (20%) | >0.9 |
| Statin Therapy, n (%) | 5 | NA | 1 (20%) | >0.9 |
| ACEi / ARB Therapy, n (%) | 5 | NA | 3 (60%) | >0.9 |
| MRA Therapy, n (%) | 5 | NA | 1 (20%) | >0.9 |
| Beta Blocker Therapy, n (%) | 6 | NA | 3 (50%) | >0.9 |
| Anticoagulation, n (%) | 5 | NA | 1 (20%) | >0.9 |
| Hemoglobin [g/dl] | 5 |  |  |  |
| Median (IQR) |  | NA | 14.60 (13.60, 15.20) |  |
| Leukocytes [n/nl] | 5 |  |  |  |
| Median (IQR) |  | NA | 6.80 (5.40, 7.50) |  |
| Thrombocytes [n/nl] | 5 |  |  |  |
| Median (IQR) |  | NA | 184 (151, 304) |  |
| C-Reactive Protein [mg/l] | 5 |  |  |  |
| Median (IQR) |  | NA | 3 (2, 3) |  |
| Creatinine [mg/dl] | 5 |  |  |  |

Deciphering human HFpEF at single cell resolution
| Characteristic | N | Control, N = 12 | HFpEF, N = 6 | p-value<sup>†</sup> |
| --- | --- | --- | --- | --- |
| Median (IQR) |  | NA | 1.15 (1.03, 1.20) |  |
| Brain Natriuretic Peptide [pg/ml] | 4 |  |  |  |
| Median (IQR) |  | NA | 610 (464, 889) |  |
| LV-EF [%] | 6 |  |  |  |
| Median (IQR) |  | NA | 54 (51, 61) |  |
| IVSDd [mm] | 6 |  |  |  |
| Median (IQR) |  | NA | 17.50 (17.00, 18.75) |  |
| E/E' | 6 |  |  |  |
| Median (IQR) |  | NA | 14.0 (12.8, 18.0) |  |
| TAPSE [mm] | 6 |  |  |  |
| Median (IQR) |  | NA | 19.50 (16.25, 22.00) |  |

<sup>†</sup> Fisher's exact test; Wilcoxon rank sum test

After quality control, a total of 4,157 nuclei were retained. We integrated the data with datasets from twelve healthy septal myocardial samples taken from [19] using canonical correlation analysis. This approach achieved sufficient integration without any obvious batch effects (**Fig 1c, Fig. S1a**). When investigating overall transcriptional differences using Pearson correlation of the aggregate expression of each gene per sample (pseudo-bulk), we found no obvious differences between HFpEF etiologies, however we found a distinct difference between control and HFpEF samples (**Fig. S1b**). Quality metrics after filtering were comparable between samples (**Fig. S1d**).

Unsupervised clustering using the Louvain algorithm returned 15 clusters (**Fig. S1c**). Semi-automated cell type annotation using the CellTypist package identified nine cardiac major cell types including cardiomyocytes, endothelial cells, fibroblasts and pericytes (**Fig. 1d**) which were confirmed using canonical cell type markers (**Fig. 1e**). In addition, macrophages defined the by expression of *LYVE1*, *MERTK* and *MSR1*, mast cells (*CPA3*, *KIT*, *HDC*) and T cells (*ITK*, *CD2*, *CD69*) were annotated (**Fig. 1d, e**).

Quantification of the annotated cell types revealed differences in cellular distribution in HFpEF compared to healthy control samples (**Fig. 1f**) with comparable distributions across samples (**Fig. S1d**). We noted a significant decline in cardiomyocytes (p < 0.001), while endothelial cells were increased (p < 0.001) in HFpEF compared to controls. Macrophages and T-cells were enriched in HFpEF patient-derived samples (p = 0.014 and 0.048, respectively).

To assess overall transcriptomic differences between cell types in HFpEF, we analyzed differentially expressed genes (DEGs) using the Model-based Analysis of Single-cell Transcriptomics (MAST) in a down-sampled data set to ascertain equal statistical power between cell types. Among the abundant cell types, the transcriptome was strongest regulated in fibroblasts, suggesting that the largest transcriptional differences occur in this cell type. In addition, other stromal cells such as endothelial cells and pericytes as well as inflammatory cells showed major differences in gene expression, more than in we observed in cardiomyocytes (**Fig. 1g**).

DGE analysis of the whole dataset revealed a striking cell type-specific alteration of gene expression (**Fig. S2a-b**). Only three genes, namely the nuclear pore complex *NPIPB4* and *NPIPB5* and the transcription factor *GABPB2* were up-regulated in all cell types (p adj < 0.05, log2FC < +1; **Fig. S2c**), a finding which was validated in a published bulk RNA sequencing data set [23] only for *GABPB2* (**Fig. S2d**). Only the long non-coding RNA *MALAT1* was commonly down-regulated (**Fig. S2c**). However, its regulation could not be confirmed in the publicly available bulk RNA-Seq dataset (**Fig. S2d**).

### HFpEF cardiomyocytes show signs of metabolic structural alterations

Next, we assessed specific alterations in cardiomyocytes. Among the down-regulated genes, we found a pronounced enrichment of genes associated with aerobic respiration, mitochondrial function and fatty acid metabolism (**Fig. 2a**). Especially genes from the REACTOME term ‘aerobic respiration and respiratory electron transport’ were consistently downregulated with little heterogeneity between samples, including different HFpEF etiologies (**Fig. 2b**). These data were consistent with, however more pronounced and less heterogeneous compared to previously published bulk RNA-sequencing data of cardiac tissue from HFpEF patients (**Fig. S3a**).

**Figure 2.**
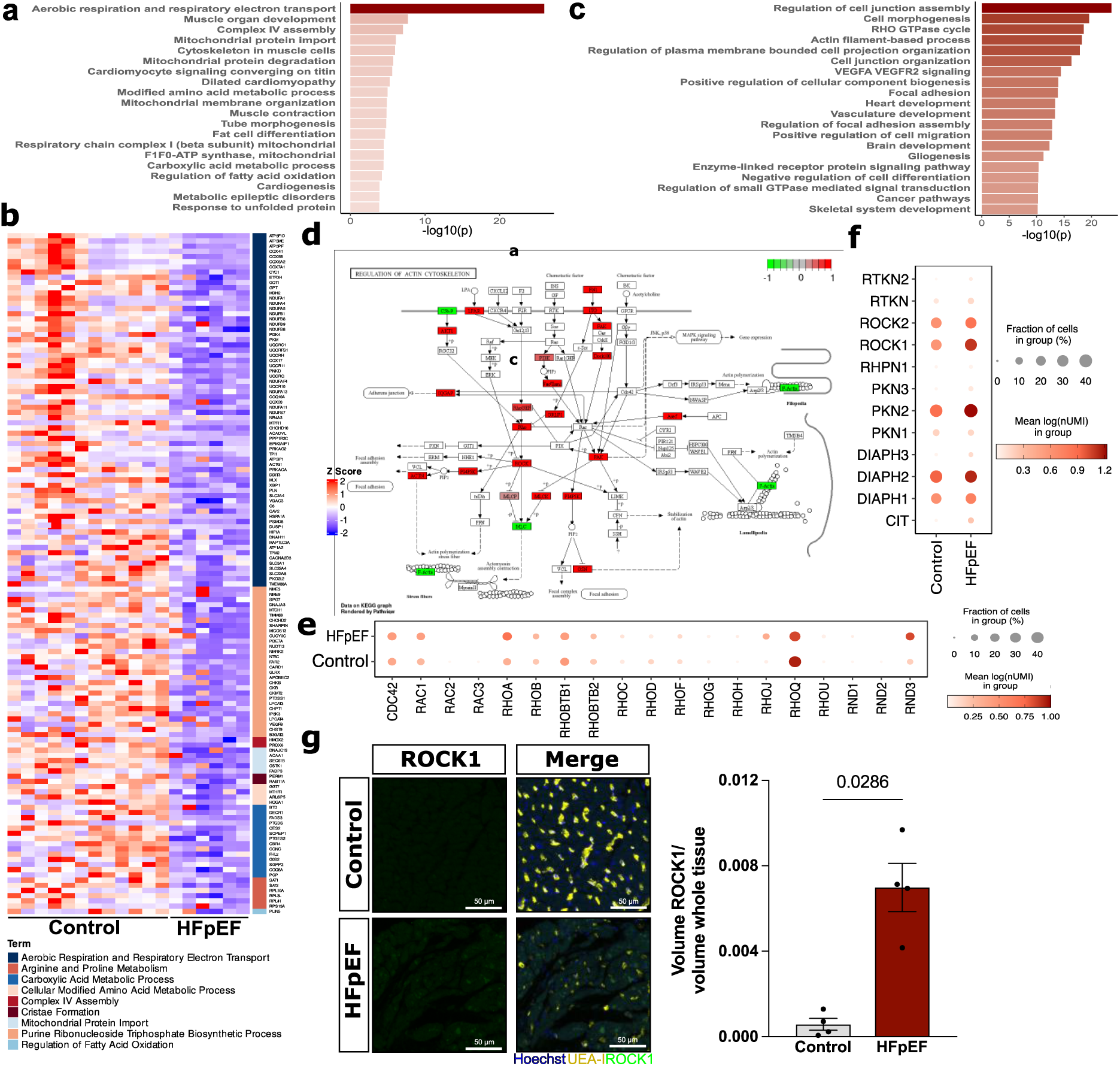
HFpEF induces structural and metabolic changes in cardiomyocytes. **a**, Metascape analysis of downregulated (adj p <0.05, log2FC < −1.0) in cardiomyocytes. **b**, Heatmap of significantly regulated genes contained in the indicated Reactome, GO and CORUM terms. **c**, Metascape analysis of upregulated DEGs (adj p < 0.05, log2FC > +1.0) in cardiomyocytes. **d**, Pathview illustration of the KEGG Pathway ‘Regulation of actin cytoskeleton’. **e**, Average expression of small GTPase genes in HFpEF and Control cardiomyocytes. **f**, Average expression of downstream effectors of small GTPases in Control and HFpEF cardiomyocytes. **g**, Representative immunofluorescence images (left) and quantification of ROCK1 signal (right) of control and HFpEF myocardial tissue stained for Hoechst, UEA-I and ROCK1 (n = 4). adj p, adjusted p value; log2FC, log2 fold-change; DEGs, Differentially expressed genes; GO, Gene ontology; CORUM, Comprehensive Resource of Mammalian Protein Complexes; KEGG, Kyoto encyclopedia of genes and genomes; HFpEF, Heart Failure with preserved Ejection Fraction; ROCK1, rho-associated coiled-coil-contaiing protein kinase 1; UEA-I, Ulex europaeus agglutinin I.

Up-regulated DEGs in cardiomyocytes were associated with the REACTOME terms “RHO GTPase cycle” and related terms on cell junction and focal adhesions as well as actin filament remodeling (**Fig. 2c**). Analysis of components of this pathway revealed that many upstream and downstream components were significantly regulated (**Fig. 2d**). Among the members of small GTPases only *RHOA* and *RND3* are significantly up-regulated (**Fig. 2e**). From the common downstream targets of these small GTPases, only *ROCK1* was significantly upregulated (**Fig. 2f**). KEGG pathway analysis suggested integrin-mediated induction of Rho-signaling with a downstream induction of *ROCK1* and subsequently *ACTN*, but a downregulation of F-Actin (**Fig. 2d**). This might be indicative of cardiomyocyte relaxation dysfunction and hypertrophy, as commonly observed in HFpEF [16]. Immunofluorescence stainings for ROCK1 in myocardial section from HFpEF patients revealed a robust upregulation which was primarily localized to cardiomyocytes (**Fig. 2g**).

In addition, we detected a high enrichment of genes associated with VEGF signaling and vascular development (**Fig. 2b**). These include members of the VEGF family (**Fig. S3b**), such as VEGFA, which is induced by 2.5-fold and may be involved in a pro-angiogenic adaptive response to hypoxia.

### HFpEF endothelial cells show increased signs of apoptosis and fetal gene reactivation

Coronary microvascular dysfunction and capillary rarefaction is increasingly recognized as a significant factor in HFpEF [12]. Accordingly, we found increased apoptosis effector genes, such as *CASP3*, *CASP7* and *CAPN2* in endothelial cells (ECs) from HFpEF samples (**Fig. 3a**). To identify potential upstream regulators that might induce EC apoptosis, we performed ligand-receptor interaction analyses using the CellChat package. Notably, the strongest altered predicted incoming ligand-receptor interaction in HFpEF ECs was SEMA3, comprising the semaphorin 3 group of ligands and respective receptors (**Fig. 3b**). As these ligands have previously been associated with anti-angiogenic effects, such as inhibition of endothelial cell migration and induction of apoptosis, we further followed this path [36]. While no canonical downstream target genes of semaphorin 3 are known, investigating the receptors of this group revealed an upregulation of *NRP1*, *NRP2*, *PLXNA4*, *PLXND1* and a mild induction of *ROBO1* (**Fig. 3c**). This expression pattern might point towards a role for SEMA3A and SEMA3C, as they primarily signal through these receptors [37]. Accordingly, we found a downregulation of genes associated with VEGFA-signaling (**Fig. 3d**), which has previously been shown to be partially counteracted by SEMA3A and induction of genes associated with negative regulation of cell migration (**Fig. 3e**). Among the upregulated DEGs in HFpEF endothelial cells, we found many to be associated with fetal gene programs, indicated by an enrichment of the GO terms ‘tube morphogenesis’, ‘embryonic morphogenesis’ and ‘heart development’ (**Fig. 3e**). To obtain insights into the extent of the upregulated fetal genes, we identified shared upregulated genes in HFpEF ECs and a publicly available bulk RNA-sequencing dataset of fetal and adult healthy myocardium [34] (both adj p < 0.05 and log2FC > +1). Interestingly, we observed that out of 592 up-regulated genes in ECs, 127 were shared with genes upregulated in fetal hearts, accounting for 21.5% of total DEGs in HFpEF ECs (**Fig. 3f**). These genes were associated with embryonic Gene Ontology (GO)-terms, ‘such as neuron projection morphogenesis’, ‘embryonic morphogenesis’, as well as ‘extracellular matrix organization’, as previously described in [34] (**Fig. 3g**).

**Figure 3.**
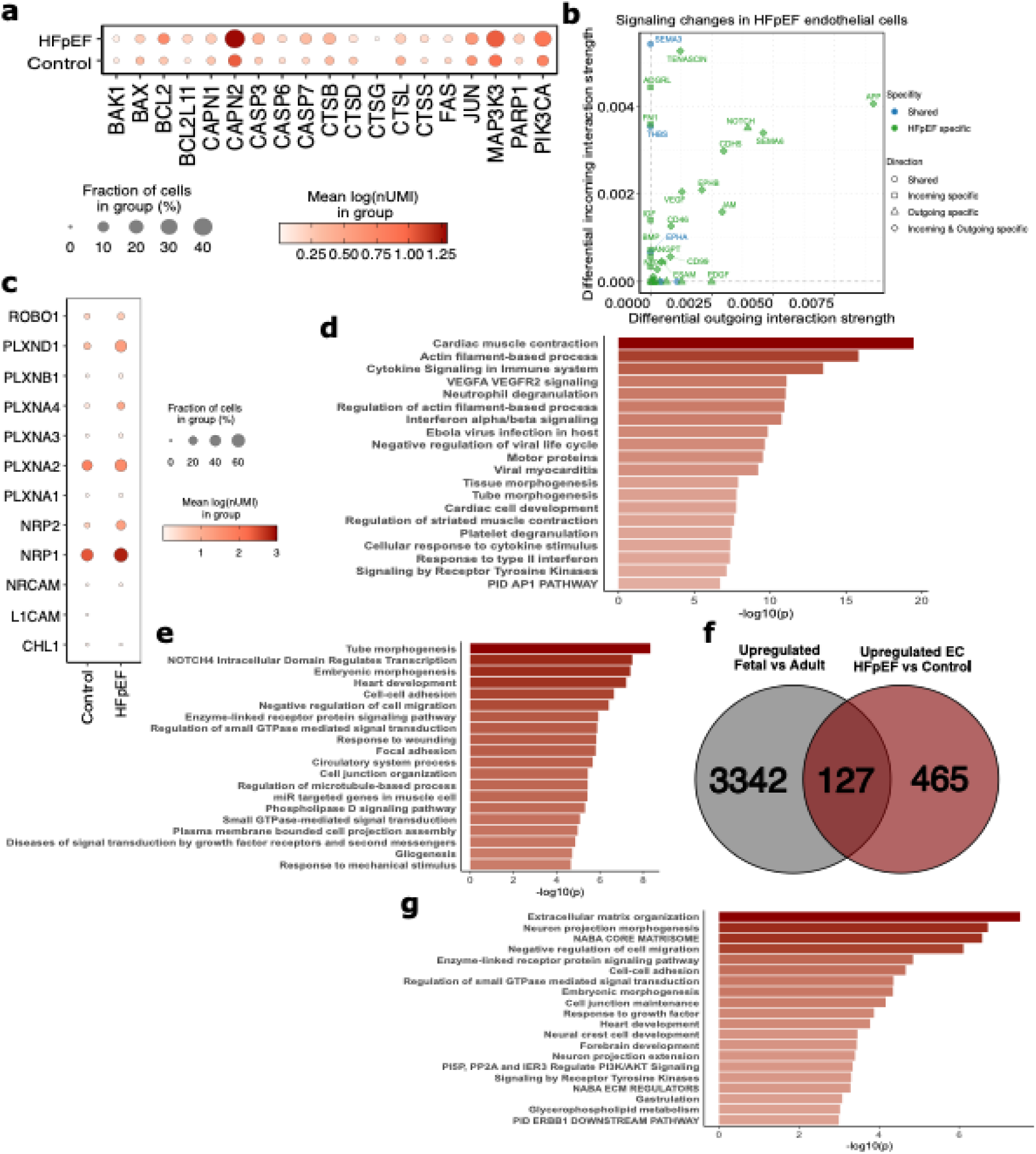
HFpEF associates with endothelial cell apoptosis and fetal gene reactivation. **a**, Average expression of apoptosis effector genes in Control and HFpEF endothelial cells. **b**, Relative changes in incoming and outgoing receptor-ligand interactions of HFpEF versus Control endothelial cells as indicated by CellChat analyses. **c**, Average expression of Semaphorin 3 receptor genes in Control and HFpEF endothelial cells. **d-e** Metascape analysis results of (**d**) downregulated (adj p < 0.05, log2FC < −1) and (**e**) upregulated DEGs (adj p < 0.05, log2FC < −1) as calculated by MAST of HFpEF versus Control endothelial cells. **f**, Venn diagram showing the number of upregulated DEGs (adj p < 0.05, log2FC > +1) as calculated by Wald test of (left) of healthy fetal versus adult myocardium from bulk RNA-sequencing and upregulated DEGs of HFpEF versus control endothelial cells (adj p < 0.05, log2FC > +1) as indicated by MAST (right). **g**, Metascape analysis results of the overlap of upregulated DEGs in fetal versus adult healthy hearts and HFpEF versus control endothelial cells. HFpEF, Heart Failure with preserved Ejection Fraction; DEGs, Differentially expressed genes; adj p, adjusted p value; log2FC, log2 fold-change; MAST, Model-based Analysis of Single-cell Transcriptomics.

### Fibroblasts in HFpEF are activated and show reduced IFNɣ signaling

Consistent with the prominent role of fibrosis in HFpEF, we observed an overall increased abundance of fibroblasts (FB; **Fig. 1f**). Using cluster-based cell-type annotation with CellTypist on fibroblasts, we did not observe an increased abundance of activated FB (**Fig. 4a-b, Fig. S4a**). However, when performing DGE analyses on the full FB cluster, we observed an overall up-regulation of the activation marker genes *COL1A1*, *COL1A2*, *COL3A1*, *COL5A1*, *FBLN1* and *LOXL1*, while *FBLN2* and the myofibroblast marker gene *ACTA2* were down-regulated (**Fig. 4c**), indicating fibroblast activation in HFpEF myocardium. Moreover, many components of the TGF-β signaling pathway were upregulated in HFpEF FB (**Fig. S4b**) Moreover, cell cycle scoring showed higher proportions of HFpEF fibroblasts to be in cell cycle phases G2/M and S, potentially indicating higher proliferative rate (**Fig. 4d**).

**Figure 4.**
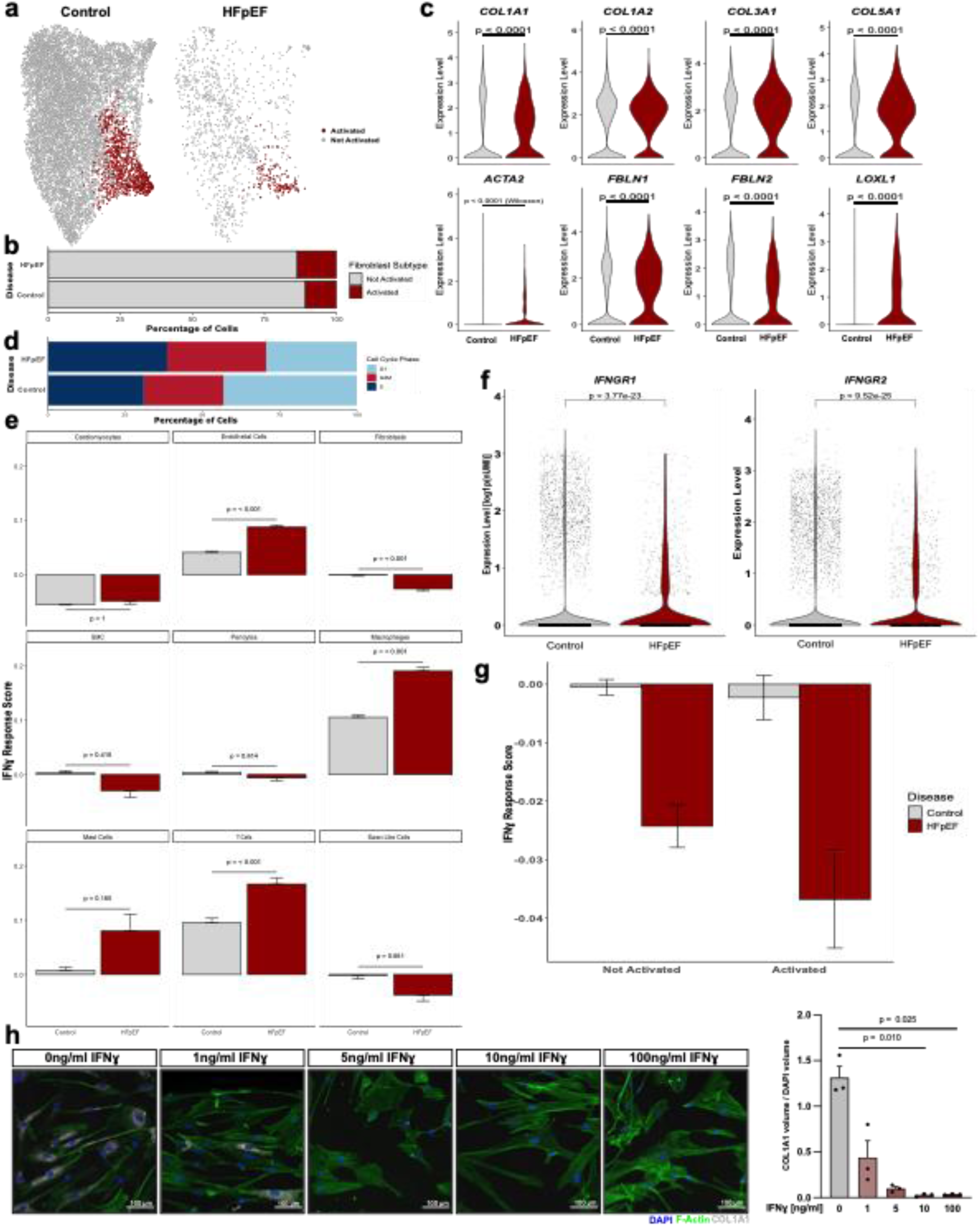
Cardiac Fibroblasts from HFpEF hearts show signs of activation and reduced IFNɣ signaling. **a,** UMAP over all fibrobla sts showing sub-populations of fibroblasts in Control and HFpEF conditions. **b**, Quantification of activated or non-activated fibroblasts after subsampling. **c**, Expression of fibroblast activation markers across all fibroblasts from Control and HFpEF samples. **d**, Frequency of predicted cell cycle phases according to cell cycle scoring in control and HFpEF fibroblasts. **e**, Bar graphs showing IFNɣ response scores per cell type for control and HFpEF fibroblasts, calculated by the average expression of IFNɣ response genes subtracted by the aggregate expression of a random control gene set per nucleus. **f**, Expression of *IFNGR1* and *IFNGR2* in control and HFpEF fibroblasts. **g**, Average IFNɣ response score in activated and non-activated fibroblasts in control and HFpEF samples. **h**, Immunofluorescence images of primary human cardiac fibroblasts treated with different concentrations of recombinant human IFNɣ stained for DAPI, F-actin and collagen 1A1 with the according quantification of collagen 1A1 volume normalized to DAPI volume (n = 3). UMAP, Uniform Manifold Approximation and Projection, HFpEF, Heart Failure with preserved Ejection Fraction; IFNɣ, interferon gamma; IFNGR1, interferon gamma receptor 1; IFNGR2, interferon gamma receptor 2; DAPI, 4’,6-Diamidin-2-phenylindol.

Most interestingly, we found a downregulation of downstream targets of interferon gamma (IFNɣ), which have previously been described in [33] specifically in HFpEF fibroblasts, whereas they were unaltered in other cell types (i.e. cardiomyocytes, smooth muscle cells or pericytes) or upregulated (T-cells, Macrophages and Endothelial cells) (**Fig. 4e**). A response score was calculated. In line with a reduced IFNɣ response score, we found a down-regulation of both IFNɣ receptor genes *IFNGR1* and *IFNGR2* in FBs from HFpEF myocardium (**Fig. 4f**). To gain further insights into a potential connection between IFNɣ signaling and fibroblast activation, we investigated the IFNɣ response score in activated and non-activated fibroblasts from control and HFpEF samples and found a reduced response score in activated FB, which was most pronounced in HFpEF samples (**Fig. 4g**). To validate these findings, we treated primary human cardiac fibroblasts with recombinant human IFNɣ *in vitro* and observed a dose-dependent reduction in collagen 1A1 protein content (**Fig. 4h**).

To investigate potential sources of IFNɣ, we created a score for IFNɣ production as described in [33]. When comparing cell types, we observed the strongest baseline score in T-cells with a further induction in HFpEF. Slight increases were observed in cardiomyocytes and pericytes (**Fig. S4c**). These data suggest a heart-intrinsic source of IFNɣ as part of an immune cell – fibroblast interaction.

### HFpEF macrophages show a pro-inflammatory phenotype and upregulation of MHC-II molecules

We observed an overall increased abundance of macrophages in HFpEF compared to control myocardium (**Fig. 1f**) with high transcriptomic changes as indicated by the number of DEGs (**Fig. 1g**). After annotation of bone marrow-derived and tissue resident macrophages within the macrophage cluster (**Fig. 5a**), we did not observe changes in the distribution of these populations between control and HFpEF (**Fig. 5b**). An analysis of the whole macrophage cluster showed a high induction of genes associated with antigen processing and inflammation signatures (**Fig. 5c**). The latter include transcriptomic signatures associated with NF-ϰB activation, such as an up-regulation of *TRAF3*, *TRAF5*, *TRAF6*, *NOD2*, *CHUK* and *IKBKB*, while the NF-ϰB inhibitor *IKBIA* was down-regulated (**Fig. S5a**). Interestingly, the transcriptomic signatures observed in HFpEF macrophages point towards increased expression of antigen presentation response, indicated by GO-terms such as ‘antigen processing and presentation of exogenous peptide antigen’ and ‘regulation of immune effector process’ (**Fig. 5c**). Accordingly, we found an upregulation of several MHC-II molecule encoding genes in HFpEF (**Fig. 5d**), indicating increased activity of the adaptive immune system. Of the nine human protein-coding genes for classical MHC-II molecules, seven were up-regulated in HFpEF macrophages. Accordingly, genes associated with MHC-II molecule transcription, i.e. *CIITA* and *CREB1*, and processing, i.e. *CD74,* were upregulated (**Fig. 5d-e**). To obtain insights into the functional associations of HLA upregulation, we combined the significantly regulated classical HLA genes (*HLA-DRA, HLA-DRB1, HLA-DRB5, HLA-DQA1, HLA-DQB1, HLA-DPA1, HLA-DPB1*) into a score, which was calculated for all macrophages. After grouping cells into quartiles based on the HLA expression score, we observed a strong enrichment of HFpEF macrophages in the upper quartiles (**Fig. 5f**). DGE analysis of the fourth (high scoring) versus the first quartile (low scoring) revealed an upregulation of genes associated with pro-inflammatory mechanisms, including GO-terms such as ‘positive regulation of cytokine production’ and ‘regulation of lymphocyte activation’ (**Fig. 5g**).

**Figure 5.**
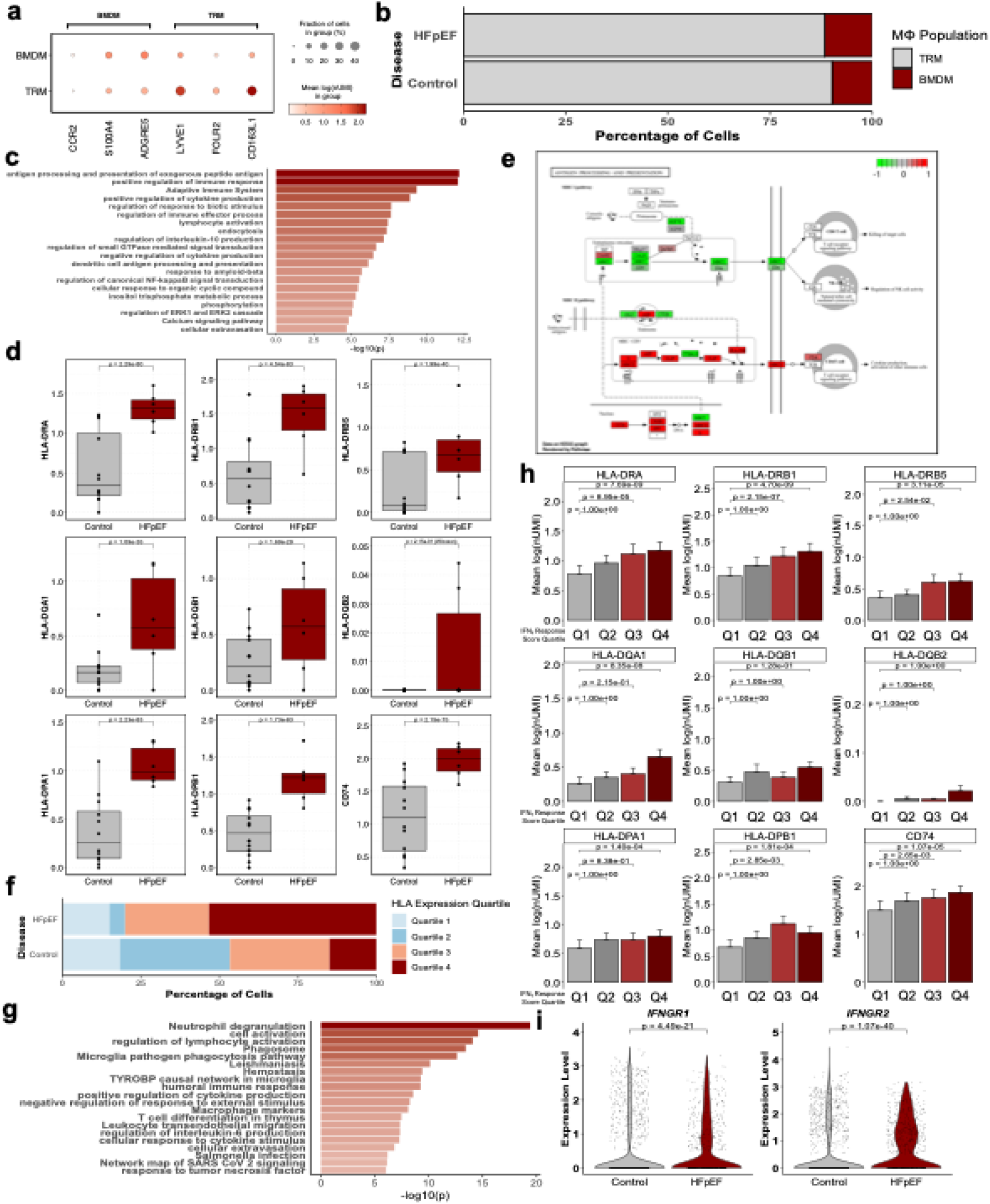
HFpEF associates with increased HLA-expression, IFNɣ-response and pro-inflammatory phenotypes in myocardial macrophages. **a**, Average expression of marker genes across macrophages sub-populations (BMDM and TRM). **b**, Frequency distribution of BMDM and TRM in Control and HFpEF samples. **c**, Metascape analysis of upregulated DEGs (adj p < 0.05, log2FC > +1) as determined by MAST in macrophages of HFpEF versus Control samples. **d**, Expression of MHC-II molecule encoding genes and *CD74* in macrophages from control and HFpEF samples. **e**, Pathview illustration of the KEGG pathway ‘Antigen Processing and Presentation’ based on the log2FC of DEGs (p adj < 0.05) of HFpEF versus control macrophages as determined by MAST. **f**, Frequency of HLA-score quartiles in macrophages from HFpEF and control myocardium. HLA-expression score was calculated by subtracting the expression of a random gene set from the aggregate expression of MHC-II encoding genes per nucleus and split in quartiles across the whole macrophage population. **g**, Metascape analysis results using upregulated DEGs (adj p < 0.05, log2FC > +1) of high versus low-HLA-scoring (4th vs 1st quartile) macrophages in HFpEF myocardium. **h**, Average expression of MHC-II molecule encoding genes and CD74 according to the quartile of the IFNɣ response score calculated by the average expression of IFNɣ target genes subtracted by the average expression of random other genes per nucleus. **i**, Expression of *IFNGR1* and *IFNGR2* in macrophages from control and HFpEF myocardium. HFpEF, Heart Failure with preserved Ejection Fraction; BMDM, Bone Marrow Derived Macrophage; TRM, Tissue Resident Macrophage; DEGs, Differentially Expressed Genes; adj p, adjusted p value; log2FC, log2 fold-change; MAST, MAST, Model-based Analysis of Single-cell Transcriptomics; MHC-II, Major histocompatibility complex class II; CD74, Cluster of Differentiation 74; KEGG, Kyoto Encyclopedia of Genes and Genomes; HLA, human leukocyte antigen; IFNɣ, interferon gamma; IFNGR1, interferon gamma receptor 1; IFNGR2 interferon gamma receptor 2.

As IFNɣ is a canonical upstream inducer of MHC-II expression is [38] and we had already observed an increased IFNɣ response score in macrophages (**Fig. 4e**), we investigated a potential association between these observations. We grouped macrophages in quartiles based on their IFNɣ response score and analyzed the expression of the significantly upregulated HLA genes between the quartiles. Indeed, we observed a higher expression for *HLA-DRA*, *HLA-DRB*, *HLA-DRB5*, *HLA-DQA1*, *HLA-DPA1* and *HLA-DPB1*, as well as for *CD74* when comparing the highest against the lowest quartile (**Fig. 5h**). In line with a role for IFNɣ signaling in HFpEF macrophages, we observed an increased expression of both IFNɣ receptors *IFNGR1* and *IFNGR2* (**Fig. 5i**).

## DISCUSSION

The data from this study provide novel insights into the transcriptional changes occurring in the heart of HFpEF patients at single-cell resolution. The most significant alterations were observed in stromal cells, which exhibited a high number of DEGs affecting multiple mechanisms and signaling pathways. In contrast, cardiomyocytes displayed modest transcriptional alterations. We found potential regulating and targetable factors of known hallmarks of HFpEF, such as RhoA/ROCK and actin cytoskeleton remodeling in cardiomyocytes. Already in 1999, it was shown that overexpression of RhoA resulted in early lethality and dilated cardiomyopathy [39]. While subsequent studies confirmed that RhoA critically regulates cardiac hypertrophy presumably via induction of hypertension, other studies also showed cardioprotective effects in acute settings [40] and pharmacologic intervention in this pathway has been discussed for more than 10 years [41]. Its contribution to HFpEF and potential implications in increased cardiomyocyte stiffness and thus diastolic dysfunction have not been studied. Furthermore, our finding of impaired fatty acid oxidation and mitochondrial dysfunction is consistent with the recently reported impaired fatty acid metabolism in hearts of HFpEF patients [9]. Interestingly, and in contrast to other cell types, in cardiomyocytes we observed signs of increased VEGFA signaling, which has been involved in altered cardiomyocyte metabolism towards increased mitochondrial biosynthesis, potentially representing a compensatory mechanism [42].

Another hallmark of HFpEF is capillary rarefaction and endothelial dysfunction, although the regulating factors are poorly understood. In this study, we observed an overall increase in endothelial cells in HFpEF. This is likely due to sampling differences, as endomyocardial biopsies from HFpEF patients are enriched in endothelial cells compared to the septal tissue samples used in other human studies. Here, we observed an increased expression of apoptosis effector genes, which associated with signs of increased Semaphorin 3 signaling. SEMA3 molecules are bona fide inducers of endothelial cell apoptosis and inhibitors of EC migration and proliferation [36, 43]. Moreover, SEMA3A has been implicated in the aging-associated loss of cardiac innervation, which resulted in increased fibrosis and diastolic dysfunction [44]. We observed an increase in the expression of fetal genes primarily in endothelial cells. While this transcriptional fetal reprogramming has been previously described in heart failure, it was mainly attributed to fibroblasts and cardiomyocytes [34, 45]. The inducing factors and functional consequences remain largely elusive, however, in endothelial cells, it is tempting to speculate that it represents a compensatory mechanism for the loss of vascular cells and subsequent hypoxia.

Among stromal cells, the most profound transcriptional changes were identified in fibroblasts. Interestingly, we did not observe an increased abundance of cells annotated as activated fibroblasts, however observed an overall increase in activation markers and signs of increased fibroblast proliferation. While this might represent a technical issue in clustering and cell type annotation, as the overall transcriptomic differences might not be large enough to be captured by clustering algorithms, it might also indicate an activation state, primarily characterized by increased ECM production and not classical myofibroblast activation. This is also supported by a downregulation of the canonical myofibroblast marker *ACTA2* in HFpEF fibroblasts. Our finding of fibroblast activation aligns with the high degree of fibrosis observed in HFpEF patients via MRI imaging [13]. Interestingly, single-cell RNA sequencing of cardiac tissue in HFpEF mouse models — induced by NO-synthase inhibition and a high-fat diet — revealed less fibroblast activation. This discrepancy suggests that these models may represent an earlier disease state or lack the pro-fibrotic component observed in human HFpEF [14].

One of the key observations in fibroblasts was a reduction in interferon gamma (IFNɣ) signaling and an association with increased fibroblast activation. IFNɣ is a cytokine with critical roles in immune regulation and antiviral defense. It is mainly produced by immune cells such as T-cells and natural killer cells, mediates pro-inflammatory responses and has been implicated in the pathogenesis of various inflammatory and fibrotic diseases. This suggests that IFNγ may contribute to the inflammatory environment driving HFpEF pathophysiology.

IFNɣ has repeatedly been shown to reduce fibroblast activation *in vitro* [46, 47]*. In vivo*, global ablation of IFNɣ induced diastolic dysfunction and a trend towards more fibrosis in a model of aldosterone infusion, uninephrectomy and high salt intake [48]. Here, we extend the *in vitro* data to human cardiac fibroblasts. The finding may have different underlying causes. On the one hand, we observed a downregulation of both genes encoding for IFNɣ receptors (*IFNGR1* and *IFNGR2*). On the other hand, we observed a strong induction of intracellular TGFβ-induced signaling components in fibroblasts. In other conditions, such as wound healing, the signaling pathways induced by IFNɣ and TGFβ have been shown to reciprocally inhibit each other, a mechanisms which is potentially contributing to the observed effects in our data [49].

In line with previous data, we found signs of increased IFNɣ production in HFpEF T-Cells [33]. This data suggests a heart-intrinsic source of IFNɣ and underlines the central role of chronic inflammation in HFpEF. In macrophages, we observed a pronounced pro-inflammatory transcriptomic signature, with a robust induction of MHC class II expression. This induction suggests an activation of adaptive immunity, highlights the potential role of macrophages in shaping the immune microenvironment in HFpEF and underscores the need for further research into the crosstalk between macrophages and adaptive immune cells in this condition. IFNɣ is an inducer of MHC class II gene expression and HFpEF macrophages showed increased transcriptomic signs of a IFNɣ response [38]. This might hint towards a pro-inflammatory polarization of macrophages through IFNɣ. However, the role of MHC-II^high^ macrophages in HFpEF and potential implications of interferons needs further investigation.

Overall, we observed an increase in macrophages under HFpEF conditions, consistent with findings in animal models and HFpEF patients with comorbidities such as obesity or metabolic syndrome [15, 50]. Using semi-supervised cell type annotation, we did not see an increase in cells annotated as bone marrow-derived macrophages. However, this analysis is limited by the lack of *CCR2* expression in our dataset, but the reduced expression of *LYVE1* and other tissue resident macrophage marker genes, as well as the increased expression of other bone marrow-derived macrophages markers, might support this finding. This finding requires further validation using other quantitative methods. In combination with an increase in overall macrophage abundance, these findings can either be caused by phenotypic switching of bone marrow-derived macrophages with a loss of CCR2 or an expansion of tissue resident macrophages. These macrophages, derived from embryonic progenitors and maintained locally through self-renewal, are highly adapted to their tissue-specific environment and perform essential homeostatic functions such as debris clearance, tissue repair, and immune regulation. However, despite their reparative roles, we observed a shift towards a pro-inflammatory phenotype, where tissue-resident macrophages in HFpEF showed profound gene expression alterations, most notably an increase in MHC class II expression. The upregulation of MHC class II genes suggests that these macrophages may acquire a pro-inflammatory function.

## LIMITATIONS

Our study is limited by the relatively low number of samples, the heterogeneity of HFpEF etiologies, and the potential bias this may introduce. As such, the study should be viewed as hypothesis-generating. While some findings have been validated, the functional consequences of the observed changes and the generalizability of our findings warrant further investigation.

## ABBREVIATIONS

HFpEF: Heart Failure with preserved Ejection Fraction
SGLT2: Sodium/glucose cotransporter 2
ATTR: Transthyretin amylyloidosis
ECM: Extracellular matrix
EMB: Endomyocardial biopsy
PCA: Principal component analysis
CCA: Canonical correlation analysis
UMAP: Uniform manifold approximation and projection
UMI: Unique molecular identifier
DEG: Differentially expressed genes
MAST: Model-based analysis of single-cell transcriptomics
snRNA-Seq: single-nucleus
VEGF: Vascular endothelial growth factor
rRNA: ribosomal RNA
DGE: Differential gene expression
FB: Fibroblasts
IFNɣ: interferon gamma
NF-ϰB: nuclear factor ‘kappa-light-chain-enhancer’ of activated B cells
MHC-II: Major histocompatibility complex class
HLA: Human leukocyte antigen
GO: Gene ontology
KEGG: Kyoto encyclopedia of genes and genomes
CORUM: Comprehensive Resource of Mammalian Protein Complexes
EC: Endothelial cells
NO: Nitric oxide

## ACKNOLEDGEMENT

We thank the patients who participated in this study. We thank Bianca Schumacher for excellent technical assistance.

## SOURCES OF FUNDING

This study was supported by the Dr. Rolf M. Schwiete Stiftung (Project 08/2018), the German Center for Cardiovascular Research (DZHK), the Deutsche Forschungsgemeinschaft (DFG) SFB1531 (project number 456687919, WP B01), the European Research Council (ERC-2021-ADG, GAP-101053352, Neuroheart) (all awarded to SD), (ERC-2021-ADG, GAP-101054899, CHIP-AVS, awarded to AMZ) and the German Society for Cardiology (Research Stipend awarded to LZ).

## DISCLOSURES

None.

## Notes

### Competing Interest Statement

The authors have declared no competing interest.

## REFERENCES

1. Mishra, S. and D.A. Kass, Cellular and molecular pathobiology of heart failure with preserved ejection fraction. Nat Rev Cardiol, 2021. 18(6): p. 400–423.

2. Schiattarella, G.G., et al., Immunometabolic Mechanisms of Heart Failure with Preserved Ejection Fraction. Nat Cardiovasc Res, 2022. 1(3): p. 211–222.

3. Roh, J., et al., Heart Failure With Preserved Ejection Fraction: Heterogeneous Syndrome, Diverse Preclinical Models. Circ Res, 2022. 130(12): p. 1906–1925.

4. Anker, S.D., et al., Empagliflozin in Heart Failure with a Preserved Ejection Fraction. N Engl J Med, 2021. 385(16): p. 1451–1461.

5. Olivotto, I., et al., Genetic causes of heart failure with preserved ejection fraction: emerging pharmacological treatments. Eur Heart J, 2023. 44(8): p. 656–667.

6. Griffin, J.M., H. Rosenblum, and M.S. Maurer, Pathophysiology and Therapeutic Approaches to Cardiac Amyloidosis. Circ Res, 2021. 128(10): p. 1554–1575.

7. Withaar, C., et al., Heart failure with preserved ejection fraction in humans and mice: embracing clinical complexity in mouse models. Eur Heart J, 2021. 42(43): p. 4420–4430.

8. Koleini, N., et al., Landscape of glycolytic metabolites and their regulating proteins in myocardium from human heart failure with preserved ejection fraction. Eur J Heart Fail, 2024. 26(9): p. 1941–1951.

9. Hahn, V.S., et al., Myocardial Metabolomics of Human Heart Failure With Preserved Ejection Fraction. Circulation, 2023. 147(15): p. 1147–1161.

10. Paulus, W.J. and C. Tschope, A novel paradigm for heart failure with preserved ejection fraction: comorbidities drive myocardial dysfunction and remodeling through coronary microvascular endothelial inflammation. J Am Coll Cardiol, 2013. 62(4): p. 263–71.

11. Luxan, G. and S. Dimmeler, The vasculature: a therapeutic target in heart failure? Cardiovasc Res, 2022. 118(1): p. 53–64.

12. Velollari, O., et al., Focusing on microvascular function in heart failure with preserved ejection fraction. Heart Fail Rev, 2025.

13. Garg, P., et al., Left ventricular fibrosis and hypertrophy are associated with mortality in heart failure with preserved ejection fraction. Sci Rep, 2021. 11(1): p. 617.

14. Lanzer, J.D., et al., Single-cell transcriptomics reveal distinctive patterns of fibroblast activation in heart failure with preserved ejection fraction. Basic Res Cardiol, 2024. 119(6): p. 1001–1028.

15. Hahn, V.S., et al., Endomyocardial Biopsy Characterization of Heart Failure With Preserved Ejection Fraction and Prevalence of Cardiac Amyloidosis. JACC Heart Fail, 2020. 8(9): p. 712–724.

16. Borbely, A., et al., Cardiomyocyte stiffness in diastolic heart failure. Circulation, 2005. 111(6): p. 774–81.

17. Daou, D., T.G. Gillette, and J.A. Hill, Inflammatory Mechanisms in Heart Failure with Preserved Ejection Fraction. Physiology (Bethesda), 2023. 38(5): p. 0.

18. Chia, Y.C., et al., Interleukin 6 and Development of Heart Failure With Preserved Ejection Fraction in the General Population. J Am Heart Assoc, 2021. 10(11): p. e018549.

19. Litvinukova, M., et al., Cells of the adult human heart. Nature, 2020. 588(7838): p. 466–472.

20. Reichart, D., et al., Pathogenic variants damage cell composition and single cell transcription in cardiomyopathies. Science, 2022. 377(6606): p. eabo1984.

21. Chaffin, M., et al., Single-nucleus profiling of human dilated and hypertrophic cardiomyopathy. Nature, 2022. 608(7921): p. 174–180.

22. Nicin, L., et al., A human cell atlas of the pressure-induced hypertrophic heart. Nature Cardiovascular Research, 2022. 1(2): p. 174–185.

23. Hahn, V.S., et al., Myocardial Gene Expression Signatures in Human Heart Failure With Preserved Ejection Fraction. Circulation, 2021. 143(2): p. 120–134.

24. Heidenreich, P.A., et al., 2022 AHA/ACC/HFSA Guideline for the Management of Heart Failure: Executive Summary: A Report of the American College of Cardiology/American Heart Association Joint Committee on Clinical Practice Guidelines. J Am Coll Cardiol, 2022. 79(17): p. 1757–1780.

25. McDonagh, T.A., et al., 2021 ESC Guidelines for the diagnosis and treatment of acute and chronic heart failure. Eur Heart J, 2021. 42(36): p. 3599–3726.

26. Fleming, S.J., et al., Unsupervised removal of systematic background noise from droplet-based single-cell experiments using CellBender. Nat Methods, 2023. 20(9): p. 1323–1335.

27. Hao, Y., et al., Dictionary learning for integrative, multimodal and scalable single-cell analysis. Nat Biotechnol, 2024. 42(2): p. 293–304.

28. Dominguez Conde, C., et al., Cross-tissue immune cell analysis reveals tissue-specific features in humans. Science, 2022. 376(6594): p. eabl5197.

29. Finak, G., et al., MAST: a flexible statistical framework for assessing transcriptional changes and characterizing heterogeneity in single-cell RNA sequencing data. Genome Biol, 2015. 16: p. 278.

30. Zhou, Y., et al., Metascape provides a biologist-oriented resource for the analysis of systems-level datasets. Nat Commun, 2019. 10(1): p. 1523.

31. Jin, S., et al., Inference and analysis of cell-cell communication using CellChat. Nat Commun, 2021. 12(1): p. 1088.

32. Tirosh, I., et al., Dissecting the multicellular ecosystem of metastatic melanoma by single-cell RNA-seq. Science, 2016. 352(6282): p. 189–96.

33. Ashour, D., et al., An interferon gamma response signature links myocardial aging and immunosenescence. Cardiovasc Res, 2023. 119(14): p. 2458–2468.

34. Spurrell, C.H., et al., Genome-wide fetalization of enhancer architecture in heart disease. Cell Rep, 2022. 40(12): p. 111400.

35. Shumliakivska, M., et al., DNMT3A clonal hematopoiesis-driver mutations induce cardiac fibrosis by paracrine activation of fibroblasts. Nat Commun, 2024. 15(1): p. 606.

36. Sakurai, A., C.L. Doci, and J.S. Gutkind, Semaphorin signaling in angiogenesis, lymphangiogenesis and cancer. Cell Res, 2012. 22(1): p. 23–32.

37. Sharma, A., J. Verhaagen, and A.R. Harvey, Receptor complexes for each of the Class 3 Semaphorins. Front Cell Neurosci, 2012. 6: p. 28.

38. Basham, T.Y. and T.C. Merigan, Recombinant interferon-gamma increases HLA-DR synthesis and expression. J Immunol, 1983. 130(4): p. 1492–4.

39. Sah, V.P., et al., Cardiac-specific overexpression of RhoA results in sinus and atrioventricular nodal dysfunction and contractile failure. J Clin Invest, 1999. 103(12): p. 1627–34.

40. Kilian, L.S., et al., RhoA: a dubious molecule in cardiac pathophysiology. J Biomed Sci, 2021. 28(1): p. 33.

41. Surma, M., L. Wei, and J. Shi, Rho kinase as a therapeutic target in cardiovascular disease. Future Cardiol, 2011. 7(5): p. 657–71.

42. Braile, M., et al., VEGF-A in Cardiomyocytes and Heart Diseases. Int J Mol Sci, 2020. 21(15).

43. Guttmann-Raviv, N., et al., Semaphorin-3A and semaphorin-3F work together to repel endothelial cells and to inhibit their survival by induction of apoptosis. J Biol Chem, 2007. 282(36): p. 26294–305.

44. Wagner, J.U.G., et al., Aging impairs the neurovascular interface in the heart. Science, 2023. 381(6660): p. 897–906.

45. Dirkx, E., P.A. da Costa Martins, and L.J. De Windt, Regulation of fetal gene expression in heart failure. Biochim Biophys Acta, 2013. 1832(12): p. 2414–24.

46. Diaz, A. and S.A. Jimenez, Interferon-gamma regulates collagen and fibronectin gene expression by transcriptional and post-transcriptional mechanisms. Int J Biochem Cell Biol, 1997. 29(1): p. 251–60.

47. Serpier, H., et al., Antagonistic effects of interferon-gamma and interleukin-4 on fibroblast cultures. J Invest Dermatol, 1997. 109(2): p. 158–62.

48. Garcia, A.G., et al., Interferon-gamma ablation exacerbates myocardial hypertrophy in diastolic heart failure. Am J Physiol Heart Circ Physiol, 2012. 303(5): p. H587–96.

49. Ishida, Y., et al., The essential involvement of cross-talk between IFN-gamma and TGF-beta in the skin wound-healing process. J Immunol, 2004. 172(3): p. 1848–55.

50. Liu, H., et al., Inflammatory Macrophage Interleukin-1beta Mediates High-Fat Diet-Induced Heart Failure With Preserved Ejection Fraction. JACC Basic Transl Sci, 2023. 8(2): p. 174–185.

